# Integrated gene-catalog and genome-resolved metagenomics reveal taxonomic and metabolic differences in the gut microbiome in irritable bowel syndrome

**DOI:** 10.64898/2026.09.05.749602

**Authors:** Mina Hojat Ansari, Mehregan Ebrahimi, Neda Khalili Sabet, Kamran Bagheri Lankarani

## Abstract

**Background:** Irritable bowel syndrome is a common gastrointestinal disorder with heterogeneous symptoms and incompletely understood pathophysiology. Gut microbiome dysbiosis has been repeatedly associated with IBS, but the extent to which taxonomic changes are accompanied by gene-level and genome-resolved differences in microbial functional potential and inferred microbial community structure remains unclear.

**Results:** We conducted a case-control shotgun metagenomic study of stool samples from patients with IBS and healthy controls. Taxonomic profiling identified distinct IBS associated compositional patterns, including enrichment of *Klebsiella pneumoniae* and *Streptococcus parasanguinis* and depletion of SCFA-associated taxa such as *Phascolarctobacterium* and *Faecalibacterium prausnitzii*. Overall microbial community composition differed between IBS and healthy controls. Assembly-based analysis generated a cohort-specific catalog of 934,495 nonredundant microbial genes. Functional profiling identified differences in carbohydrate-active enzyme families involved in dietary and host-derived glycan metabolism. Genome-resolved analysis recovered 154 nonredundant high-quality metagenome-assembled genomes and identified distinct taxonomic patterns, including HC associated *Bifidobacterium longum* and IBS associated *Dialister invisus, CAG-177 sp003538135*, and CAG-568 *sp000434395*. Metabolic analysis of the recovered genomes further revealed IBS associated enrichment of pathways related to fatty acid biosynthesis and elongation, de novo purine biosynthesis, and histidine degradation. Together, these findings demonstrate that IBS-associated microbiome differences extend beyond community composition to gene-level and genome-resolved functional potential.

**Conclusions:** This study provides a multilayered view of the IBS gut microbiome by integrating read-based taxonomic profiling, a cohort-specific microbial gene catalog, functional annotation, genome-resolved reconstruction, and metabolic-module analysis. IBS was characterized by depletion of SCFA-associated taxa, enrichment of facultative anaerobic and opportunistic taxa, altered representation of carbohydrate-active enzyme families, and distinct genome-resolved metabolic patterns. The convergence of taxonomic, gene-catalog, and MAG-level findings highlights coordinated differences in microbial composition and encoded metabolic potential in IBS and identifies candidate microbial populations and functions for validation in larger cohorts and targeted mechanistic studies.

## Background

Irritable bowel syndrome (IBS) is a common gastrointestinal disorder that affects approximately one in ten people worldwide [1]. IBS is characterized by chronic abdominal pain, bloating, and irregular bowel habits. The condition significantly affects patients’ quality of life and imposes a major socioeconomic burden through increased healthcare costs [1]. The exact pathophysiology of IBS remains unclear, but several factors have been implicated, including disturbances in gut motility, visceral hypersensitivity, involvement of the immune system, and psychological stress [2]. Over the past decade, newly developed sequencing technologies have improved our understanding of the relationship between symbiotic microorganisms that live on and within us and their contribution to health and disease [3]. Emerging evidence increasingly implicates gut microbiome dysbiosis as an important component of IBS pathophysiology. IBS has been associated with shifts in specific microbial taxa and alterations in microbial metabolic potential [4-6]. However, the biological significance of these changes and relationship to the heterogeneous symptoms of IBS remain areas of active research.

Shotgun metagenomic sequencing has transformed the study of the gut microbiome by enabling researchers to move beyond traditional 16S rRNA profiling and resolve microbial species and strains, functional genes, and encoded metabolic pathways at substantially higher resolution [7]. Unlike marker-gene sequencing, shotgun metagenomics enables direct profiling of the taxonomic and functional potential of microbial communities, providing a framework for identifying microbial features that may contribute to IBS pathophysiology. Despite increasing evidence of microbial dysbiosis in IBS, several critical questions remain unresolved. Studies have reported bacterial alteration, but inconsistencies across cohorts, clinical subtypes, sequencing methods, and analytical approaches, together with limited high-resolution functional data, have made it difficult to define a reproducible biological signature of IBS. It also remains unclear how microbial functional potential differs among IBS subtypes and whether these differences are associated with their heterogeneous clinical features. Because many previous studies have relied on amplicon sequencing, strain-level variation and the connection between specific microbial populations and their encoded functions remain incompletely characterized. Reconstruction of metagenome-assembled genomes (MAGs) can overcome several of these limitations by linking community-level changes to genome-resolved microbial populations and their metabolic capacities. Moreover, whether IBS associated taxonomic and functional shifts are accompanied by broader reorganization of bacterial, archaeal, viral, and eukaryotic community associations remains poorly understood. Integrated genome-resolved and cross-domain analyses therefore offer an opportunity to move beyond descriptive taxonomic dysbiosis, but their application in IBS remains limited.

To address these gaps, in the current study, we performed shotgun metagenomic sequencing of stool samples from IBS patients and healthy controls to identify key microbial taxa associated with IBS and its subtypes, and determine whether IBS associated dysbiosis extends beyond taxonomic composition. We integrated species and strain-level taxonomic profiling with construction and functional annotation of a nonredundant microbial gene catalog, reconstruction of metagenome-assembled genomes, analysis of encoded metabolic modules, and profiling of antibiotic-resistance and virulence-associated features. We further applied cross-domain co-occurrence network analysis to examine whether compositional and functional shifts were accompanied by changes in the inferred organization of bacterial, archaeal, viral, and eukaryotic communities. This multi-layer framework allowed us to identify IBS associated taxa and genome-resolved populations, investigate subtype-linked patterns, and characterize coordinated changes across microbial composition, encoded metabolic potential, and community organization.

## Methods

### Study Population and Experimental Design

Patients with irritable bowel syndrome were invited to participate in the current study while attending the outpatient clinic of Shiraz University of Medical Science. All the patients met the Rome IV criteria [8] for diagnosis of IBS. Then they were evaluated by a gastroenterologist prior to enrollment to alternative diagnoses and confirm the diagnosis. The IBS patients were classified into subtypes based on Bristol Stool Form scale characteristics following the Rome IV criteria for subcategorizing IBS patients: IBS with constipation (IBS-C), IBS with diarrhea (IBS-D), and IBS with mixed habits (IBS-M) [8]. The healthy controls (HC) had no history of chronic or recurring gastrointestinal symptoms. The main inclusion criteria included 20 to 50 years of age, no history of GI tract surgery (excluding appendectomy or cholecystectomy), no history of lactose malabsorption, diabetes, polyp, inflammatory bowel diseases (IBD), celiac disease, any chronic disease including chronic kidney, chronic obstructive pulmonary and collagen vascular disease. Participants were also excluded if they were pregnant, had a history of any diagnosis that might disrupt the gut microbiota, had consumed prebiotics, probiotics or synbiotics during the previous two years, or had been treated with antibiotics during the three months before enrollment. In total, 180 individuals were examined, of which 42 subjects were eligible and included in the study. Verbal informed consent was obtained from all participants. The study was approved by the ethical committee of the Shiraz University of Medical Science (IR.SUMS.REC.1398.188). The cohort and shotgun metagenomic sequencing dataset analysed here were previously used in a virome-focused study [9]. The present study represents a distinct analysis of the same 42 metagenomes, focusing on read-based microbial taxonomy, gene-level functional potential, genome-resolved reconstruction, metabolic modules, and microbial association structure.

### Sampling, DNA Isolation, and Whole-Genome Sequencing

Fresh fecal samples were collected from all participants in plastic tubes on-site while attending the outpatient gastroenterology clinic at Shiraz University of Medical Science for physical examination. The collected samples were immediately transferred on ice and then stored at −80°C in the laboratory. The library preparation was performed as previously described in [9]. Briefly, DNA was extracted using the NucleoSpin® Tissue DNA isolation kit (Genomic DNA from tissue, Macherey-Nagel, Germany) from 250 mg of feces following the manufacturer’s protocols. Shotgun metagenomic sequencing libraries were prepared using the Nextera XT DNA Sample Prep Kit (Illumina Inc. San Diego, CA) and sequenced on an Illumina NextSeq500 platform with a 2 × 150 paired-end read length at the Faghihi Medical Genetics Center (FMG; Shiraz, Fars, Iran).

### Short Reads Assessment and Taxonomic Profiling

All read preprocessing and read-based taxonomic analyses were performed using versioned tools and workflow within the Galaxy platform on the European Galaxy server, (www.usegalaxy.eu) [10, 11]. Raw sequencing reads were processed to obtain high-quality clean reads for further analysis. Reads were first quality-filtered and adapter sequences were removed using Trim Galore v0.6.6 [12]. Human-derived reads were removed by aligning the sequences to the human reference genome (GRCh38.p13) using BBMap [13]. Then, taxonomic profiling was conducted using Kraken2, Bracken, and MetaPhlAn through the initial version of the *“*Metagenomic Taxonomy analysis*”* workflow implemented in Galaxy [10] and publicly available online (https://usegalaxy.eu/published/workflow?id=7491883694fff308). Kraken2 was used for taxonomic classification, followed by Bracken for abundance estimation, while MetaPhlAn provided complementary microbial profiles for cross-method comparison.

### Metagenomic Assembly and Construction of the Gene Catalog

Metagenomic assembly, gene prediction, and sequence clustering for gene-catalog construction were performed using versioned tools implemented within Galaxy [10]. The clean reads of each sample were individually assembled for gene catalog construction using MetaSPAdes [14]. Assembled contigs were used for gene prediction with Prodigal [15] using metagenomic mode (-p meta). Genes shorter than 100 bp were discarded, and the remaining genes were clustered (≥ 95% identity and ≥ 90% overlap) using MMseqs2 [16]. The final nonredundant gene catalog was subjected to taxonomic and functional assignment, in which gene-level taxonomy was predicted through alignment to the Uniprot TrEMBL database [17]. KEGG (Kyoto Encyclopedia of Genes and Genomes) [18] annotation were assigned using KOBAS v3.0.3 [19]. The NCBI Clusters of Orthologous Groups (COGs) annotation results were extracted with eggNOG-mapper v2.0.1b [20], and carbohydrate-active enzymes (CAZymes) were annotated with the dbCAN database using HMMER [21]. The protein domains were annotated using Pfam database. Antibiotic resistance genes (ARGs) and virulence factors were identified using PathoFact v1.0 [22]. To estimate gene abundances, clean reads from each sample were aligned to the gene catalog using BWA-MEM [23], and read counts were computed using Samsum (https://github.com/hallamlab/samsum). Normalization was performed using fragments per kilobase of gene sequence per million mapped reads [24]. For functional comparisons, the abundance of genes assigned to the same functional annotation were aggregated within each sample before statistical comparison between study groups.

### Genome Reconstruction

Gut microbial genomes from healthy controls and patients with IBS were reconstructed using a group-wise co-assembly and multiple binning tools implemented within the Galaxy platform [10]. Briefly, MetaSPAdes software [14] was used to co-assemble clean reads from each study group (IBS-C, IBS-D, IBS-M, and HC), generating a total of four co-assemblies. Co-assembling samples within groups was intended to improve binning of low-abundance populations while still avoiding the overloading of the assembler with excessive diversity from a co-assembly of all samples [25]. Metagenome-assembled genomes (MAGs) were recovered using three binning algorithms: MetaBAT2 [26], MaxBin2 [27], and CONCOCT [28]. Then the bins were unified into a final bin set using the metaWRAP Bin refinement module (-c 70 -x 5 options) [29]. CheckM [30] was used to estimate completeness and contamination of the final bin sets. The resulted bins were then imported into the anvi’o workflow [31]. Genome completeness and redundancy were further assessed based on a set of 71 universal bacterial single-copy genes, and all bins were manually inspected to have less than 5% redundancy.

High-quality MAGs were defined as having ≥70% completeness and ≤5% contamination based on CheckM [30] assessment. The MAGs obtained from all four groups were pooled, and duplicate MAGs (identical MAGs defined independently in different study groups), were identified using an average nucleotide identity (ANI) threshold of ≥ 99% with the anvi-dereplicate-genomes workflow. Duplicate MAGs were discarded, and a final nonredundant set of MAGs was retained. The nonredundant MAGs were then assigned taxonomy using GTDB-Tk [32] with the Genomes Taxonomy Database (https://gtdb.ecogenomic.org/). Functional annotation of the MAGs was also performed within the anvi’o framework [31], using the COG, Pfam, KOfam (KEGG orthologs), and CAZyme databases.

Group specific metabolic enrichment was then calculated across the study groups. For phylogenetic analysis, the anvi’o phylogenomics workflow was used based on a set of single-copy core ribosomal proteins [33, 34]. Amino acid sequences of these marker genes were retrieved using *“*anvi-get-sequences-for-hmm-hits*”* with the return best hit, get aa sequences, and concatenate options. Multiple sequence alignment was performed using MUSCLE v3.8.1551 [35], and phylogenomic trees were generated using *“*anvi-gen-phylogenomic-tree*”* [31].

### Haplotyping

The number of haplotypes represented by each MAG was estimated using the anvi’o command *“*anvi-gen-variability-profile*”* [31], and DESMAN [36], following procedures described in detail in previous research [37]. The anvi’o command *“*anvi-gen-variability-profile*”*, with the *“*quince mode*”* option was used to export single nucleotide variant (SNV) information for all MAGs. Then the variations in core genes (in this case, SNVs in a set of 71 core genes that having a single copy in all known bacteria and archaea), and their co-occurrence across samples were analyzed using DESMAN, which estimates the number of haplotypes represented by each MAG. DESMAN requires some prior information regarding the haplotype exception to estimate the potential number of haplotypes in the population. Therefore, each MAG was decomposed into 1, 2, 3, 4, 5, 6, 8, 10, and 12 candidate haplotypes, with 500 iterations performed for each model in four replicate runs using different random seeds. Mean posterior deviance was plotted for each MAG and candidate haplotype number. Finally, the optimal number of haplotypes for each MAG was selected based on SNV uncertainty of less than %10 and posterior deviance was at least 10% [36].

### Analysis of Metabolic Modules and Enrichment

The level of completeness for a given KEGG module was assessed using the *“*anvi-estimate-metabolism*”* with default parameters. This program uses functional annotations of genes mapped to KEGG orthologs to infer metabolic module completeness. Then to identify enriched metabolic modules, we applied the *“*anvi-compute-metabolic-enrichment*”* tool, which performs enrichment analysis based on the results from *“*anvi-estimate-metabolism*”*. The analysis employs a binomial generalized linear model (GLM) to fit a logistic regression to the presence of each metabolic module across groups. Statistical significance was evaluated using a Rao test, with both uncorrected p values and Benjamini Hochberg corrected q values reported. A metabolic module was considered significantly enriched if the q value was < 0.05.

### Microbial association network analysis

Microbial association structures between HC and IBS were evaluated using species and strain level abundance profiles derived from the taxonomically annotated gene catalogue. The main feature set included all features detected in at least 50% of HC samples and at least 50% of IBS samples, irrespective of their mean abundance, retaining recurrent low-abundance taxa while excluding highly sparse features. Sensitivity analyses used shared within-group prevalence thresholds of 20%, 25%, 30%, 40%, 60%, 70%, and 80%. Zero values were replaced using a sample-wise multiplicative zero-replacement procedure, after which sample profiles were closed to proportions and centered log-ratio transformed [38]. Pearson correlations were calculated separately for HC and IBS. Group comparisons included mean absolute correlation, the root-mean-square difference between correlation matrices, and total, positive, and negative edge densities at an absolute-correlation threshold of 0.40. Species-level cross-domain associations were evaluated separately, and edge-density sensitivity was assessed at absolute-correlation thresholds of 0.35, 0.50, and 0.60.

Group differences were evaluated using 1,000 label permutations preserving the observed group sizes of 17 HC and 25 IBS samples. Permutation p values were calculated using the finite-sampling correction [39], and Benjamini-Hochberg correction was applied across the prespecified network metrics. The effect of unequal group sizes was assessed over 500 iterations by comparing all 17 HC samples with 17 IBS samples selected without replacement. Uncertainty was additionally evaluated using 500 balanced bootstrap resamples of 17 samples per group. Edge-level stability was assessed over 500 within-group bootstrap resamples. Stable edges were required to retain the same correlation sign and an absolute correlation of at least 0.40 in at least 70% of resamples, with a 95% bootstrap interval excluding zero. Additional sensitivity analyses used feature-wise half-minimum zero replacement, Spearman correlations, and comparisons between corresponding CLR–Pearson and SparCC correlation matrices [40].

As a secondary analysis, an aggregated taxon abundance matrix derived from the gene catalogue, in which each taxon was linked to its assigned KEGG functions, was evaluated using the same global network metrics. Network nodes represented taxa, while the KEGG annotations were used only to characterize their associated functional potential. The individual pathways or modules were not treated as network nodes. Subtype-specific networks were not included in inferential comparisons because the IBS-C, IBS-D, and IBS-M groups contained only 7–10 participants.

### Statistical analysis

All statistical analyses were performed with R v4.3.3 [41] and Python v3.12.2. Multiple-testing corrections was applied where appropriate using the Benjamini-Hochberg procedure, with statistical significance set at a false discovery rate (FDR) < 0.05 [42]. Alpha diversity was assessed using the Shannon and Simpson indices in phyloseq [43], with statistical differences evaluated using the Wilcoxon rank-sum and Kruskal Wallis tests, where appropriate. Beta diversity was calculated using Bray Curtis dissimilarity with the Vegan package [44] and visualized using Principal Coordinates Analysis (PCoA). Statistical significance of beta diversity differences was tested using PERMANOVA, implemented via the adonis2 function in the Vegan package, with Benjamini-Hochberg correction applied to multiple pairwise comparisons.

Differentially abundant taxa were identified using DESeq2 [45], which models count data with a negative binomial distribution and applies the Wald test. As a complementary association method, MaAsLin2 was applied to analyze relative abundance data [46]. ANCOM-BC was used as an additional compositional differential abundance method, particularly for subtype comparisons [47]. For functional analyses, abundance values for genes assigned to the same functional annotation were aggregated within each sample. Differences between HC and IBS were assessed using the Wilcoxon rank-sum test, whereas comparisons among HC and IBS-C, IBS-D, and IBS-M were assessed using the Kruskal-Wallis test. Taxonomic differential abundance results from DESeq2, MaAsLin2, and ANCOM-BC were treated as complementary analyses and reported separately. No formal consensus-selection rule was used to define candidate features. Given the small numbers of participants within the IBS-C, IBS-D, and IBS-M groups, subtype-level taxonomic and functional findings were considered exploratory and hypothesis-generating.

## Results

### Overview of Study population and design

We enrolled 42 participants, including 17 healthy volunteers and 25 IBS patients, aged 20-50 years, to investigate IBS associated gut microbiome features. The healthy subjects were free of chronic or recurrent gastrointestinal symptoms, and all participants reported no antibiotic exposure during the previous three months and no prebiotics or probiotic during the previous two years. The IBS patients were diagnosed based on the Rome IV criteria, with further confirmation by a gastroenterologist. We also divided the IBS patients into three subtypes, based on the Rome IV criteria for subcategorizing IBS patients, including: IBS-C (n = 8), IBS-D (n = 7), and IBS-M (n = 10).

Stool specimens were collected from all 42 participants and subjected to shotgun metagenomic sequencing. A description of the participants is presented in Supplementary materials Table S1. The most commonly recorded symptoms among IBS patients were abdominal pain and abdominal bloating, with bloating self-reported by almost 90% of the patients. Four patients rated bloating as their most troublesome symptom.

### Shotgun Metagenomic Sequencing and Taxonomic Profiling of Reads

Shotgun metagenomic sequencing generated a total of 0.5 billion reads across the 42 samples. Taxonomic profiling of these reads was performed using Kraken2 and MetaPhlAn (Supplementary Figure S1). Kraken2 taxonomically assigned about 50% of the reads, of which more than 97% classified as bacteria (Supplementary Figure S2). *Firmicutes* and *Actinobacteria* were the most abundant phyla in both HC and IBS, whereas *Bacteroidetes* and *Proteobacteria* were relatively more abundant in the IBS patients.

MetaPhlAn analysis detected a total of 696 species and 734 strains across all samples, with 335 species and 351 strains detected in at least 5% of samples. These species and strain level profiles provided higher taxonomic resolution for subsequent comparisons between HC and IBS, complementing the broader Kraken2/Bracken taxonomic profiles.

### Gut Microbiome Signatures in IBS and Healthy Controls

Differential abundance analysis identified taxa distinguishing IBS from HC. *Phascolarctobacterium* was significantly more abundant in HC (Figure 1A, Supplementary Figure S2). In IBS, *Klebsiella, Enterobacter, Veillonella, Rothia*, and *Haemophilus* were significantly enriched relative to HC (DESeq2, FDR q < 0.05; Figure 1B, Supplementary Figure S3).

**Figure 1.**
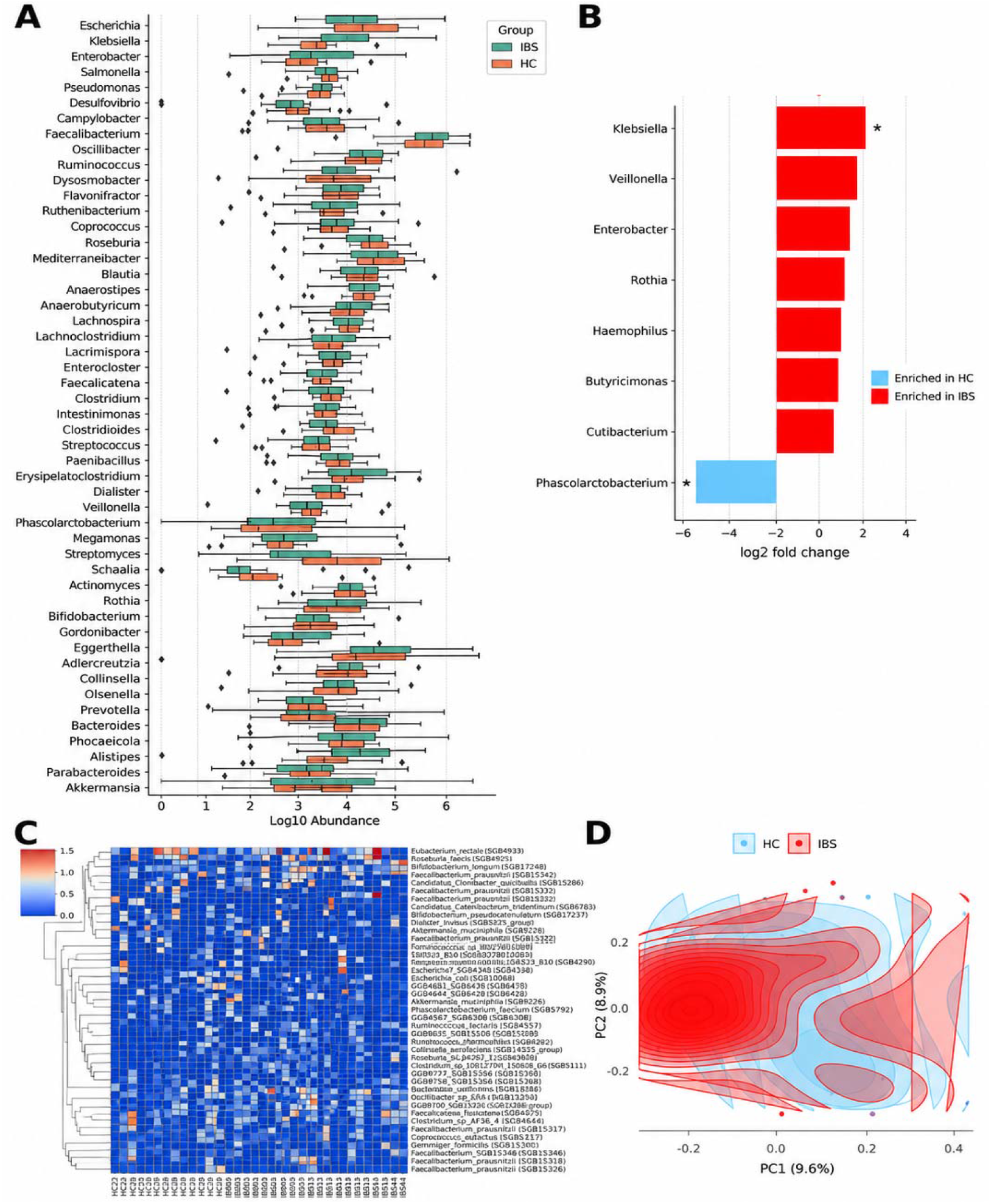
Distinct microbial signatures between IBS and HC. **A** Genus-level composition of the gut microbiome in participants with IBS (n = 25) and HC (n = 17). Boxplots show log10-transformed relative abundances of the most abundant genera, ordered by median abundance across all samples. **B** Differentially abundant genera between IBS and HC identified using DESeq2 (shown; Benjamini– Hochberg adjusted p values). Positive log2 fold change values indicate enrichment in IBS, whereas negative values indicate enrichment in HC. Genera also identified as significant by MaAslin2 are shown in bold and marked with an asterisk (*). **C** Heatmap of the 50 most abundant bacterial species and strains in IBS and HC. Abundances are shown on a log10 scale, with rows clustered by similarity and columns ordered by clinical groups. **D** Principal-coordinate analysis of Bray-Curtis dissimilarities between IBS and HC. Ellipses indicate 95% confidence regions (PERMANOVA R^2^ = 3.67%, p = 0.018).

At the species level, *Faecalibacterium prausnitzii* was one of the most prevalent species across all studied groups, being detected in almost every participant (Figure 1C). Within IBS patients, the *F. prausnitzii* strains SGB15316 and SGB15342 were detected in more than 90% of the participants, whereas *F. prausnitzii* strain SGB15333 was significantly enriched in HC, suggesting strain-level differences between IBS and HC. Taxa associated with HC included key butyrate producers such as *Holdemanella porci* and *Roseburia spp*., which were absent or significantly reduced in IBS patients (Supplementary Figure S4). In contrast, taxa associated with IBS included opportunistic species such as *K. pneumoniae, Streptococcus parasanguinis, Erysipelatoclostridium ramosum*, and *Veillonella parvula* (Supplementary Figure S4). Notably, *K. pneumoniae* (SGB10115), *Streptococcus parasanguinis* (SGB8071), and *Veillonella parvula* (SGB6939) were not only highly prevalent in IBS patients but also significantly more abundant (Supplementary Figure S4).

Furthermore, subtype-specific analyses revealed that *K. pneumoniae* (SGB10115) was significantly enriched in association with IBS-D patients. In Addition, Alistipes onderdonkii (SGB2303), *A. finegoldii* (SGB2301), *Bifidobacterium longum* (SGB17248), and *Blautia faecis* (SGB4820) were detected in nearly all IBS-D patients but were rare in HC. In IBS-C patients, *Sellimonas intestinalis* (SGB4617) was significantly enriched, along with *Agathobaculum butyriciproducens* (SGB14993), which exhibited notably high prevalence in IBS patients. Moreover, the *Coprococcus eutactus* strain SGB5118 was detected more frequently in IBS-C patients. In contrast, the *C. eutactus* strain (SGB5121) and *Streptococcus salivarius* (SGB8007) were highly prevalent in IBS-M patients. Notably, *S. salivarius* (SGB8007) was generally more prevalent in IBS patients than in HC, reinforcing its potential role in IBS pathology.

Alpha diversity did not show significant variation between IBS and HC but indicated general trends toward lower microbial diversity in IBS patients. In contrast, beta diversity analysis revealed a significant difference in gut microbial composition between IBS and HC (PERMANOVA: R^2^ = 3.67%, p = 0.018; Figure 1D). These microbial shifts suggest a transition toward a dysbiotic gut community in IBS patients.

### Gut Microbial Gene Catalog Highlights IBS-Associated Shifts

We independently assembled high-quality reads from each sample to create a gene catalog of gut microbes (Supplementary Figure S1). A ssembly of the 42 samples generated 485,583 contigs and 1.2 million genes, with an average gene length of 638bp. Of these, 492,034 genes (approximately 41%) were identified as complete. After removing genes smaller than 100 bp and clustering at 95% amino acid identity, we recovered 934,495 nonredundant human gut microbial gene (HGMG).

Among the nonredundant genes, approximately 92% (859,120 genes) were assigned to Uniprot TrEMBL, of which 818,371 genes could be taxonomically classified as bacteria (98%), archaea (0.2%), eukaryote (0.02%) and virus (0.2%) (Figure 2A). These genes were assigned to 19 phyla, 76 families, 196 genera, 3043 species, and 638 strains. The most abundant bacterial phyla were *Firmicutes* (73% of the bacteria), *Actinobacteria* (9% of the bacteria), *Bacteroidetes* (8% of the bacteria), *Proteobacteria* (3% of the bacteria), and *Verrucomicrobia* (1% of the bacteria).

**Figure 2.**
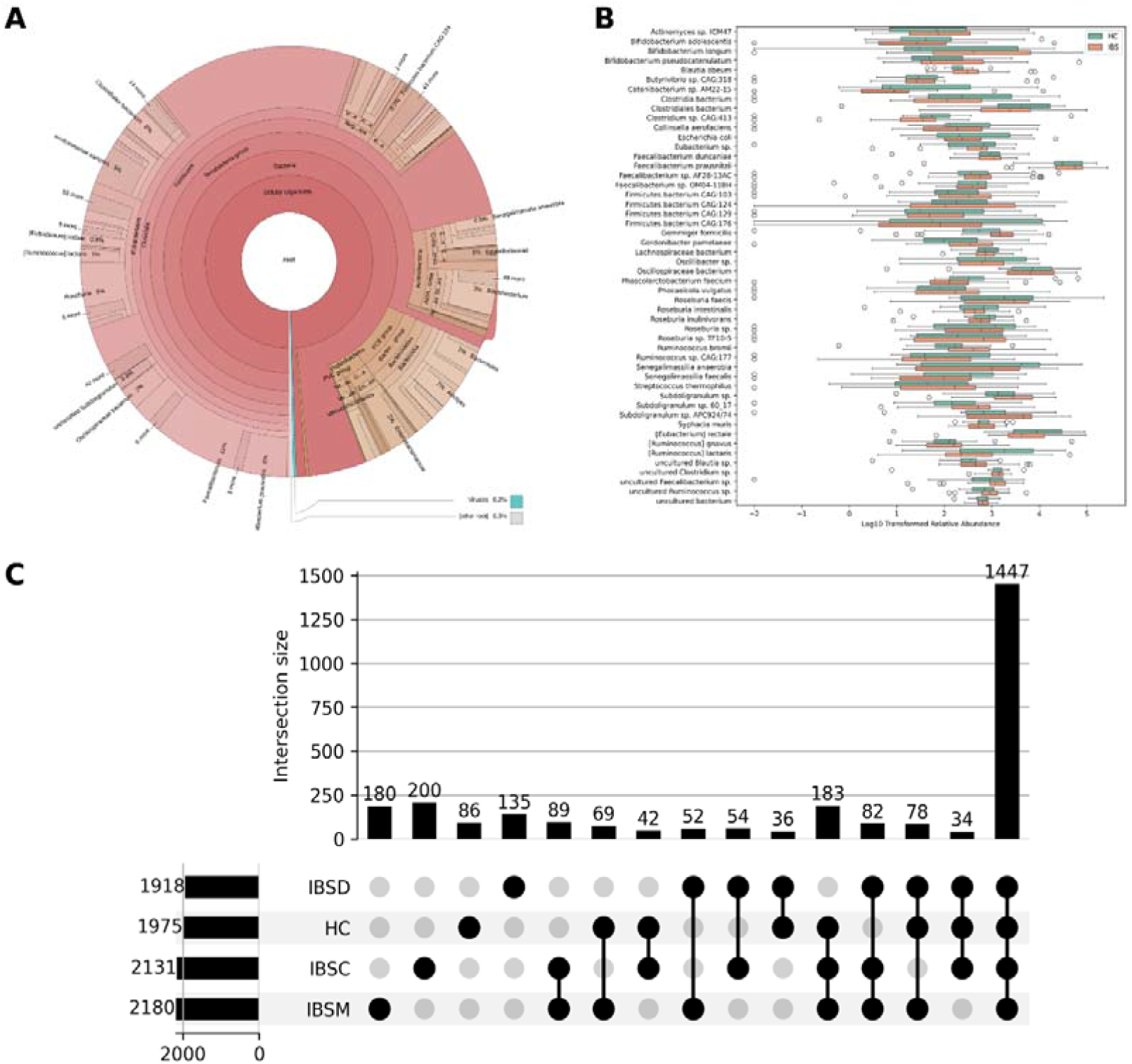
Human gut microbial gene catalog and prevalence patterns across IBS subtypes and HC. **A** Krona plot showing the taxonomic distribution of the nonredundant gene catalog recovered from 42 metagenomes. **B** Species-level composition of the core microbiome, defined as taxa present in ≥90% of all samples. Boxplots show log□□- transformed relative abundances for the 50 most abundant species among the 182 identified core species in IBS and HC. C UpSet plot shows the distribution of bacterial species across HC and IBS subtypes (IBSC, IBSD, IBSM) using a ≥10% prevalence threshold. Bars indicate the number of shared species, and connected dots indicate the groups in which each species was detected.

We further analyzed the prevalence of annotated microbiome across the samples. Taxa detected in at least 90% of all samples were defined as members of the core gut microbiome. Core microbiome analyses identified four phyla, nine families, 25 genera, 182 species, and 46 strains common across all samples (Supplementary Table S2). Among these, two phyla, five families, 12 genera, 37 species, and eight strains were detected in all 42 samples (Supplementary Table S2). The abundance of these 182 bacteria species includes more than 97% of the total abundance of 3043 annotated species, emphasizing their high prevalence and ecological importance in the human gut microbiome (Figure 2B).

To explore additional variability and potential IBS associated taxa, we applied a broad prevalence threshold to include taxa present in at least 10% of the samples. This analysis aimed to capture taxa that may not be part of the core microbiome but differed between the HC and IBS groups. In particular, we identified 16 genera and 195 species associated with IBS, appearing in at least 10% of IBS samples but being distinctly less prevalent or absent in HC (Figure 2C). These taxa may reflect IBS associated microbiome shifts and provide further insight into the potential role of the microbiome in IBS pathophysiology. Further investigation into IBS subtypes and their associated microbial gene alteration, revealed an uncultured *AlphaProteobacteria* bacterium that was uniquely prevalent in approximately 40% of IBS-C samples and absent from the other groups. Furthermore, we found that species from *Klebsiella Sp*. were significantly associated with IBS samples, particularly with IBS-D samples. These findings highlight the presence of taxa or their enrichment that, although not highly prevalent across all IBS subtypes, may serve as markers of variability or shifts within specific IBS subgroups.

### Functional Insights into the Gut Microbial Gene Catalog

The nonredundant genes were functionally annotated using the KEGG, COG, Pfam, and CAZymes databases. A total of 56,156, 638,009, and 891,805 genes were assigned to CAZyme families, COG functions, and Pfam domains, respectively. Additionally, 259,776 genes were assigned to KEGG orthologous groups, of which 50,164 genes were further assigned to KEGG pathways. These annotations enabled comparison of the functional potential encoded by the microbiomes of participants with IBS and healthy controls.

Analysis of CAZymes highlighted differences in carbohydrate-metabolism potential between HC and IBS. A total of 248 unique CAZyme families were retained for summary analysis (Supplementary Table S3), including 25 associated with HC, 26 associated with IBS, and 197 shared families showing differential abundance. HC samples showed higher abundance of glycoside hydrolase families such as GH43-23 and PL12, which are involved in the degradation of hemicellulose and pectin. In contrast, IBS samples exhibited higher abundance of families including GH43-35, GH70, and GH125, which are associated with host-derived glycan metabolism and simpler carbohydrate substrates.

Pfam analysis identified 3,758 unique domains in the summary analysis, including 326 associated with HC, 514 associated with IBS, and 2,918 shared domains showing differential abundance (Supplementary Table S4). HC-associated domains included the SHOCT domain, a conserved motif involved in bacterial signaling. In contrast, IBS samples exhibited higher abundance of the DNA polymerase III psi subunit, suggesting potential shifts in replication or stress-response processes. These results indicate functional differences in domains related to cofactor utilization, DNA replication, and redox regulation.

Complementing these findings, COG analysis identified 3,123 unique functions in the summary analysis (Supplementary Table S5), including 265 associated with HC, 474 associated with IBS, and 2,384 shared functions showing differential abundance. Prominent HC-associated functions included the predicted ATPase of the AAA+ superfamily and the lipopolysaccharide export system protein LptC, which are involved in protein quality control and membrane integrity. IBS-associated functions included the multidrug efflux-pump subunit AcrA and the conserved protein YbgA, potentially reflecting microbial stress adaptation and host interaction. Shared functions showing differential abundance included DNA polymerase DinP and ABC-type amino-acid transport systems. These findings suggest differences in microbial functions related to membrane dynamics, stress responses, and nutrient processing.

KEGG pathway analysis further revealed differences in microbial functional potential between HC and IBS. A total of 269 unique KEGG pathways were retained in the summary analysis (Supplementary Table S6). Pathways such as pyrimidine degradation and the ethylmalonyl pathway, both involved in carbon metabolism, were associated with HC. In contrast, IBS samples showed higher abundance of pathways including lipoic acid biosynthesis and the acetate kinase pathway, indicating potential differences in cofactor metabolism and short-chain fatty acid production.

To explore functional variation across IBS subtypes, KEGG pathway profiles were compared among HC, IBS-C, IBS-D, and IBS-M. A total of 279 unique pathways were retained in the subtype analysis (Supplementary Table S7). The observed subtype-associated patterns included citrate-cycle activity and gluconeogenesis in IBS-M, and heme and pyocyanine biosynthesis in IBS-D. IBS-C samples showed higher abundance of pathways including CMP-Neu5Ac and sphingosine biosynthesis. Given the small subtype sample sizes, these findings were interpreted as exploratory evidence of functional heterogeneity among IBS subtypes.

Overall, these results indicate differences in the functional potential of the microbiomes of participants with IBS and healthy controls. The observed patterns involved microbial functions related to nutrient processing, stress adaptation, cofactor metabolism, and interactions with the host.

### Microbial association analyses reveal no widespread network reorganization in IBS

The main shared-prevalence feature set retained 676 species-level features, comprising 665 bacterial, seven eukaryotic, two archaeal, and two viral taxa, together with 132 strain-level features. Using all available samples, HC showed descriptively higher species-level mean absolute correlation and edge density than IBS. However, label-permutation tests did not support differences in mean absolute correlation or edge density after multiple-testing correction (both BH-adjusted q = 0.819), and the overall correlation-matrix difference was also nonsignificant (q = 0.971). Total, positive, and negative edge densities and species-level cross-domain association metrics likewise showed no significant differences after correction. Repeated equal-size comparisons produced intervals spanning zero for species-level differences in mean absolute correlation and edge density.

Strain-level comparisons also showed no significant differences in mean absolute correlation, edge density, or overall correlation structure between HC and IBS (all BH-adjusted q ≥ 0.681). Analyses using more inclusive prevalence thresholds did not identify additional group differences. CLR-Pearson and SparCC species-level correlation matrices were concordant in HC and IBS (r = 0.852 and 0.863, respectively), indicating that the conclusion was not driven solely by the correlation estimator. The secondary analysis of 64 KEGG-annotated taxa showed descriptively stronger associations in HC, but the differences were not significant after multiple-testing correction (all q ≥ 0.206) and were sensitive to the zero-replacement procedure. Overall, the analyses did not provide robust evidence of widespread microbial-network fragmentation, altered cross-domain organization, or functional-network restructuring in IBS.

### Genome-resolved metagenome analysis revealed microbiome taxonomic indicators of IBS

We constructed the microbial genomes from the metagenomics sequences (Supplementary Figure S1). A total of 221 high-quality MAGs with ≥70% completeness and ≤5% contamination were recovered, of which 56 MAGs had ≥90% completeness and <2% contamination. After removing the redundant MAGs (sharing an average nucleotide identity of ≥ 99%), a final set of 154 nonredundant MAGs were retained. Taxonomic classification using the Genomes Taxonomy Database Toolkit (GTDB-TK), assigned the MAGs to 52 genera and 63 species from 6 bacterial phyla, including *Firmicutes, Actinobacteriota, Bacteroidota, Proteobacteria*, Verrucomicrobiota, and Cyanobacteria (Supplementary Figure S5). We then used DESMAN to investigate whether we could detect haplotypes in each MAG, which could be representative of major strain types. The 154 nonredundant MAGs comprised one to four haplotypes (Supplementary Figure S6).

Genome-resolved comparisons revealed taxonomic differences between HC and IBS, consistent with the read-based taxonomic profiles. *Bifidobacterium longum* was enriched in HC, whereas *Dialister invisus* and less characterized species, including *CAG-177 sp003538135* and CAG-568 *sp000434395*, were enriched in IBS (Figure 3A). In addition, members of the order *Oscillospirales*, particularly the genus *Ruminococcus*, were more prevalent in patients with IBS.

**Figure 3.**
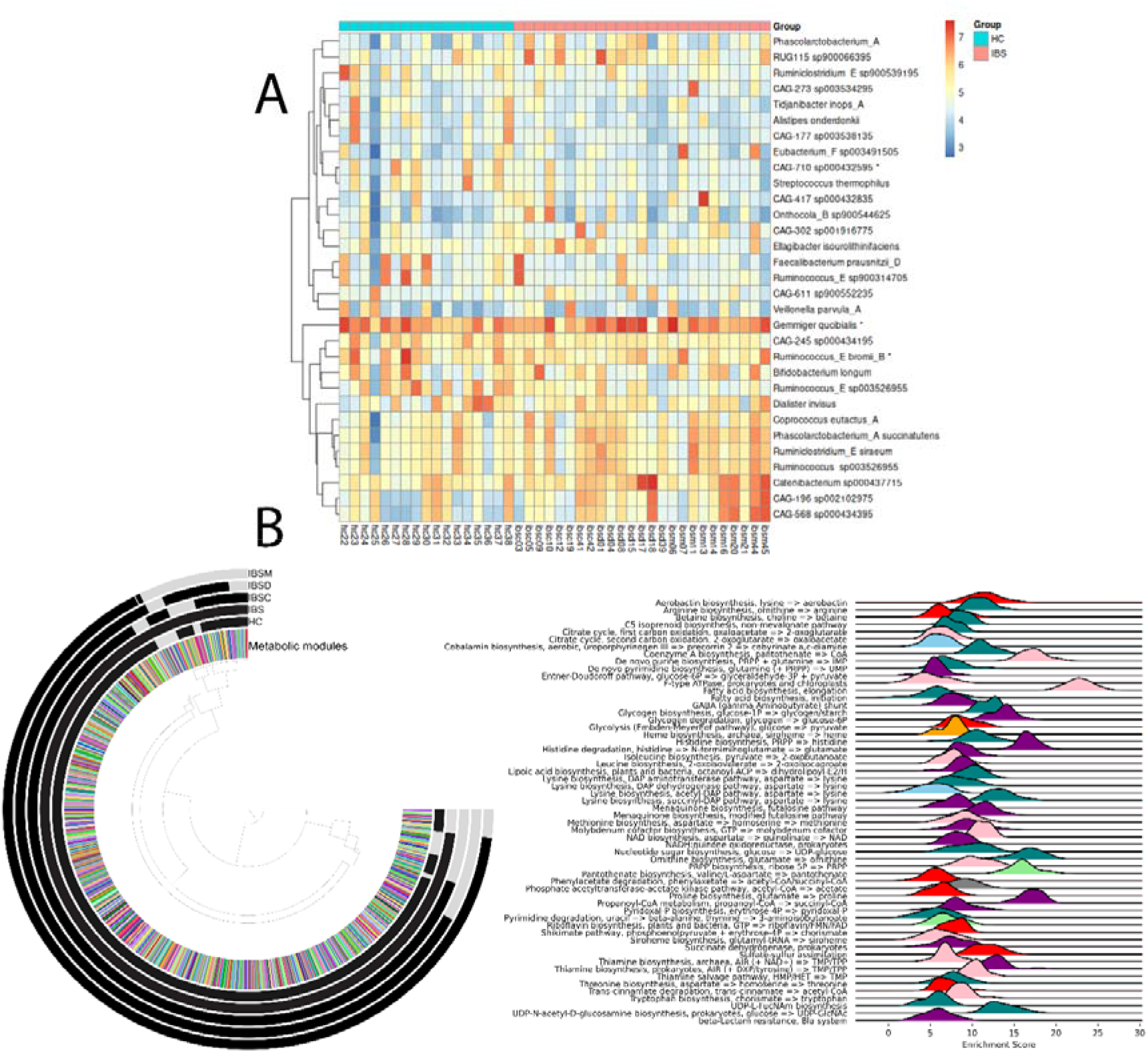
Genome-resolved metagenomics reveals taxonomic and functional differences between IBS and HC. **A** Heatmap showing species represented by MAGs that differed significantly between IBS and HC based on DESeq2 analysis. Abundances are normalized and shown on a log□□ scale: rows are clustered by similarity, and samples are grouped by clinical category. Taxa also identified as significant by ANCOM-BC are marked with asterisks (*). **B** Top: anvi’o circular display of metabolic modules showing functional variation and enrichment patterns across HC, IBS, and the IBS subtypes (IBS-C, IBS-D, IBS-M). Concentric rings represent the groups, from the innermost to the outermost ring, and shaded sectors indicate modules enriched in each group. Bottom: ridgeline plots showing enrichment-score distributions for significant modules. Colors indicate the group or combination of groups in which each module was enriched sky blue, HC; orange, IBS-C; red, IBS-D; olive, IBS-M; purple, IBS-C + IBS-D; light green, IBS-C + IBS-M; teal, IBS-D + IBS-M; and pink, all IBS subtypes.

The analysis of metabolic pathways also revealed significant differences between HC and individuals with IBS. Fatty acid biosynthesis and elongation pathways were significantly enriched in IBS, indicating increased genome-encoded potential for fatty acid metabolism that may influence motility and inflammation. Purine biosynthesis and histidine degradation were lower in HC but enriched in IBS, potentially reflecting increased histamine production, a compound linked to gut inflammation and hypersensitivity. Notably, ornithine biosynthesis, a key pathway in the urea cycle and polyamine synthesis, was predominantly enriched in IBS-D, potentially aiding epithelial repair and gut barrier function in diarrhea-predominant IBS. In contrast, the phosphate acetyltransferase-acetate kinase pathway was enriched in HC, IBS-M, and IBS-D but reduced in IBS-C, suggesting impaired microbial acetate production in IBS-C. This reduction may contribute to slower motility, disrupted microbial cross-feeding, and low-grade inflammation, distinguishing IBS-C from other IBS subtypes. Additionally, de novo purine biosynthesis was elevated across all IBS subtypes, indicating increased cellular turnover and stress responses within the gut environment. However, pathways associated with gut homeostasis, including cobalamin (vitamin B12) biosynthesis and lysine biosynthesis, were enriched in HC but suppressed in IBS, reflecting a loss of protective metabolic functions in IBS. These findings highlight specific metabolic disruptions in IBS and provide insights into potential microbial and host-microbiome interaction mechanisms underlying IBS pathophysiology (Figure 3B).

Our analysis of virulence factors (VFs) and antibiotic resistance genes (ARGs) revealed a higher overall abundance of both VFs and ARGs in IBS than in HC. The most prevalent resistance categories included multidrug, beta-lactam, aminoglycoside, and tetracycline resistance. However, statistical testing using the Wilcoxon rank-sum test did not confirm a significant difference between IBS and HC after false-discovery-rate correction. Patients with IBS tended to show a higher prevalence of resistance genes related to beta-lactam, multidrug, and fluoroquinolone resistance.

To further investigate virulence associated functions, we assessed key VF-related mechanisms, including iron uptake, bacterial adhesion, and toxin production. However, we found no significant increase in these pathways in IBS relative to HC. Similarly, no significant enrichment of adhesion-related or toxin-producing genes was observed in IBS. This suggests that, although IBS-associated microbial communities may harbor a greater load of virulence-associated bacteria, this did not translate into an increased presence of specific pathogenic pathways. Across IBS subtypes (IBS-C, IBS-D, and IBS-M), no significant differences in VFs or ARGs abundance were detected, suggesting that the observed trends were characteristic of IBS overall rather than being subtype-specific.

Overall, our findings indicate that IBS microbiomes exhibit a trend toward a greater virulence and resistance gene burden, but without significant enrichment of specific virulence pathways. Future studies integrating host microbiome interactions and functional metagenomics could provide further insight into the role of microbial functional shifts in IBS pathophysiology.

## Discussion

In the current study, we performed a high-resolution metagenomic analysis of the gut microbiome in patients with IBS compared with HC using taxonomic profiling, functional annotation, recovery of MAGs, and microbial association analysis. Our findings identified distinct microbial signatures in IBS patients, consistent with previously reported dysbiosis patterns, while uncovering novel associations and providing new insights into functional alterations. These microbial changes suggest functional shifts, particularly those linked to host glycan metabolism, gut barrier integrity, and inflammatory responses, providing insights into their potential role in IBS pathophysiology and possible therapeutic implication.

We observed that *Firmicutes, Actinobacteria, Bacteroidetes*, and *Proteobacteria* were the dominant phyla in all participants, with *Firmicutes* and *Proteobacteria* detected in 100% of the samples (Supplementary Table S2). These findings are consistent with previous studies that identified these phyla as dominant members of the human gut microbiota [48, 49]. Identification of these phyla in nearly all participants underscores their fundamental role in maintaining gut homeostasis [50]. Within the defined core microbiome, genera such as *Faecalibacterium, Bifidobacterium*, and *Bacteroides* (Supplementary Table S2), contributing to gut metabolic functions, including SCFA production, fiber degradation, and immune modulation [51]. Their consistent presence in every sample suggests that they are not only important contributors to gut health, but also part of a shared microbial backbone across individuals.

The stability of the core microbiome structure observed in our study aligns with previous reports suggesting that major bacterial phyla remain relatively conserved in IBS, while alterations occur primarily at the genus and species levels [52]. However, IBS patients exhibited specific alterations in microbial diversity and composition compared to HC. These compositional shifts may have functional consequences that contribute to IBS symptoms.

One of the most noticeable findings was the depletion of key SCFA-producing taxa in IBS patients, including the propionate producing genus *Phascolarctobacterium*, and the butyrate producing taxa *Faecalibacterium prausnitzii*, and *Roseburia spp*. Butyrate serves as an energy source for colonic epithelial cells, modulates inflammation, maintains gut barrier integrity, and regulates immune responses [53]. It is well established that butyrate has significant anti-inflammatory effects [54], and its reduction has been implicated in inflammatory bowel diseases [55]. The depletion of beneficial bacteria, including *F. prausnitzii*, and *Roseburia spp*., has been previously reported in association with IBS [56]. However, some taxa identified in previous studies were not significantly altered in our study. In addition, several taxonomic changes observed in this study, for example a reduction of *Phascolarctobacterium* in IBS patients, have not been widely reported. The significant depletion of *Phascolarctobacterium* in IBS patients suggests a loss of taxa involved in SCFA production, which is important for maintaining gut homeostasis. *Phascolarctobacterium* is a known propionate producer involved in energy metabolism and potentially in gut-brain axis interactions [57]. However, its role in IBS remains unclear because previous studies have reported inconsistent associations. For example, one study found enrichment of *Phascolarctobacterium* in diarrhea predominant IBS (IBS-D) patients [58], whereas another study linked its abundance to successful responses to a low-FODMAP diet [59]. These contrasting findings suggest that its association with IBS, may depend on additional factors, such as diet, host metabolism, and IBS subtypes, highlighting the need for further investigation.

The observed increase in *Proteobacteria*-associated taxa, including *Klebsiella* and *Enterobacter*, supports the hypothesis that gut dysbiosis in IBS is driven partly by an over-representation of facultative anaerobes and opportunistic pathogens [60]. These bacteria have been associated with gut inflammation, metabolic dysregulation, and increased gut permeability [61]. They may also contribute to increased lipopolysaccharide (LPS) biosynthesis, which can trigger immune responses and disrupt gut barrier integrity. So, contributing to gastrointestinal symptoms, particularly in IBS subtypes characterized by increased intestinal permeability [62-64].

In contrast to the depletion of beneficial SCFA-producing taxa, patients with IBS exhibited significant enrichment of facultative anaerobes and opportunistic taxa, including *Klebsiella pneumoniae, Enterobacter, Veillonella, Rothia*, and *Haemophilus*. These taxa have previously been associated with intestinal inflammation, metabolic dysregulation, and altered gut permeability [65]. Among them, *K. pneumoniae* emerged as a distinguishing microbial feature of IBS, particularly in patients with IBS-D, in whom it was significantly enriched. *Klebsiella Sp*ecies are known for their ability to metabolize host-derived glycans, produce lipopolysaccharides (LPS), and evade immune responses, which may contribute to intestinal inflammation and gut barrier dysfunction [66].

The association of *Klebsiella pneumoniae* with IBS symptoms, including bloating and stool irregularities, further supports the hypothesis that specific members of the gut microbiota may contribute to symptom severity through metabolic and immunomodulatory pathways. Previous studies have linked increased *Klebsiella* abundance to gut dysbiosis in gastrointestinal disorders, including inflammatory bowel disease, where it has been implicated in low-grade inflammation and epithelial barrier disruption [67]. The increased prevalence of *Klebsiella* in patients with IBS in our study may also be related to its role in bile acid metabolism, as previous studies indicate that *Proteobacteria*, including *Klebsiella*, can influence bile acid transformation and gut motility [68].

Beyond *Klebsiella*, other enriched taxa, such as *Veillonella parvula* and *Rothia* mucilaginosa, are known for their ability to thrive in an oxygen limited but inflammation-driven gut environment. *Veillonella* species have been associated with lactate fermentation, converting lactate into short-chain fatty acids such as acetate and propionate, which may contribute to increased gut fermentation and gas production [69, 70], potentially exacerbating bloating and discomfort in patients with IBS. Similarly, *Rothia* species, which are commonly found in the oral and respiratory microbiomes, have been linked to gastrointestinal dysbiosis in IBS and other gut disorders, suggesting possible cross-seeding from the oral cavity to the gut [71].

In addition, the observed enrichment of *Enterobacter* and *Haemophilus* in IBS suggests an expansion of facultative anaerobes that can thrive under dysbiotic gut conditions. Previous studies have similarly reported increased abundance of *Enterobacter* species in patients with IBS [72]. This shift in facultative anaerobes composition may contribute to inflammation and oxidative stress within the gut [73], although further research is needed to clarify the potential roles of *Enterobacter* and *Haemophilus* in IBS pathophysiology. Our findings indicate that patients with IBS have a higher prevalence of facultative anaerobes and opportunistic bacteria, which may contribute to gut dysbiosis by promoting intestinal permeability, inflammation, and altered metabolic processes. Although it remains unclear whether these microbial shifts are a cause or consequence of IBS, their association with IBS symptomatology suggests that targeting these microbial signatures through microbiome based interventions may represent a potential therapeutic strategy.

These taxonomic shifts were accompanied by differences in microbial functional potential between IBS and HC, as revealed by functional metagenomic analysis. The depletion of butyrate-producing bacteria may further affect epithelial barrier integrity, as butyrate plays a crucial role in maintaining barrier function and regulating gut inflammation [53]. Consistent with these metabolic alterations, we also identified distinct shifts in microbial functional potential related to carbohydrate metabolism. HC microbiomes exhibited higher abundance of GH43-23 and PL12, which are involved in the degradation of hemicellulose and pectin. In contrast, IBS microbiomes showed higher abundance of GH43-35, GH70, and GH125, suggesting differences in carbohydrate and host-glycan processing. These findings reinforce the idea that IBS associated dysbiosis involves not only taxonomic shifts but also potential functional adaptation of the microbiome to the altered gut environment. Additionally, our KEGG pathway analysis revealed differences in microbial metabolic potential. HC microbiomes were enriched in pyrimidine degradation and the ethylmalonyl pathway, whereas IBS microbiomes showed higher abundance of lipoic acid biosynthesis and the acetate kinase pathway. These findings suggest differences in carbon metabolism, cofactor metabolism, and short-chain fatty acid production between HC and IBS.

Microbial association analysis did not identify consistent evidence of widespread network reorganization in IBS. Although HC showed descriptively higher species-level correlation strength and edge density, these differences were not supported after label-permutation testing, equal-size resampling, bootstrap assessment, or sensitivity analyses across prevalence and correlation thresholds. Species-level cross-domain associations, strain-level networks, and the KEGG annotated taxon network likewise showed no robust group-level differences. Thus, while subtle or taxon-specific ecological changes cannot be excluded, the present cohort does not support widespread microbial-network fragmentation or functional-network restructuring in IBS. Larger cohorts will be needed to determine whether more modest alterations in microbial association structure occur in IBS.

Furthermore, genome-resolved metagenomic analysis revealed IBS-associated differences in microbial populations and their encoded metabolic potential. The increased abundance of species such as *Dialister invisus* and *CAG-177 sp003538135* suggests potential roles for these taxa in IBS associated dysbiosis. Metabolic-module enrichment analysis identified fatty acid biosynthesis, purine biosynthesis, and histidine degradation as enriched in IBS, supporting differences in microbial metabolic capacity. The physiological consequences of these metabolic differences remain to be established. In addition to these metabolic adaptations, our analysis of virulence factors and antibiotic resistance genes revealed a trend toward higher abundance of these genes in patients with IBS, though the differences were not statistically significant after FDR correction. While this suggests a possible link between IBS microbiomes and increased microbial resistance capacity, further validation is needed. This trend aligns with previous studies that reported an enrichment of antimicrobial resistance genes in dysbiotic gut microbiomes, particularly under conditions associated with low-grade inflammation and altered immune responses [74]. However, despite the presence of more opportunistic bacteria in IBS, we did not observe a significant increase in classical virulence pathways such as yersiniabactin or enterobactin. Rather than being driven by direct enrichment of known virulence factors, IBS associated dysbiosis may therefore be related more to shifts in microbial composition and metabolic potential than to an increased presence of specific pathogenic traits.

By integrating read-based taxonomic profiling, construction of a cohort-specific gene catalog, genome-resolved reconstruction, metabolic-module analysis, and microbial association testing, this study provides a multilayered characterization of IBS-associated microbiome variation. The cohort size, particularly within the IBS subtype groups, and the cross-sectional design mean that subtype-specific patterns and causal interpretations should be considered exploratory. Subtype-associated differences may also partly reflect variation in stool consistency and intestinal transit, while detailed dietary information was not available for adjustment. Moreover, the metagenomic findings represent encoded functional potential rather than directly measured microbial activity, and the gene-catalog functional comparisons were based on nominal significance. Nevertheless, the convergence of taxonomic, gene catalog, and genome-resolved findings identifies coherent IBS-associated patterns, particularly the depletion of SCFA-associated taxa and differences in carbohydrate-utilization and metabolic potential. These findings provide a focused foundation for validation in larger longitudinal cohorts and for targeted functional investigation of the microbial processes potentially involved in IBS.

## Conclusions

By integrating read-based taxonomic profiling, a cohort-specific gene catalog, genome-resolved reconstruction, and metabolic-module analysis, this shotgun metagenomic study identifies convergent differences in microbial composition and encoded functional potential between IBS and healthy controls. IBS was characterized by depletion of SCFA-associated taxa, enrichment of facultative anaerobic and opportunistic taxa, differences in carbohydrate-active enzyme families involved in dietary and host-derived glycan metabolism, and distinct metabolic signatures among IBS associated MAGs. Notably, microbial association analyses did not support widespread network reorganization, indicating that the IBS associated differences detected in this cohort were more evident in targeted taxonomic and functional shifts than in global microbial co-occurrence structure. Together, these multilayered findings identify candidate microbial and functional features for validation in larger independent cohorts and provide a foundation for mechanistic studies aimed at clarifying the role of the gut microbiome in IBS.

## List of abbreviations

ARGs: Antibiotic resistance genes
CAZymes: Carbohydrate-active enzymes
COGs: Clusters of Orthologous Groups
ENA: European Nucleotide Archive
FDR: False discovery rate
HC: Healthy controls
IBS: Irritable bowel syndrome
IBS-C: IBS with predominant constipation
IBS-D: IBS with predominant diarrhea
IBS-M: IBS with mixed bowel habits
KEGG: Kyoto Encyclopedia of Genes and Genomes
MAGs: Metagenome-assembled genomes
SCFAs: Short-chain fatty acids
SNV: Single-nucleotide variant
VFs: Virulence factors

## Declarations

### Ethics approval and consent to participate

This study was approved by the Ethics Committee of Shiraz University of Medical Sciences (approval no: IR.SUMS.REC.1398.188). All participants were informed about the study procedures and their right to withdraw at any time and provided verbal informed consent before participation.

### Availability of data

The raw metagenomic sequencing data generated in this study are available in the European Nucleotide Archive (ENA) under project accession PRJEB104707, with sample accession IDs ERS28186854–ERS28186934. The corresponding metadata are provided in the supplementary tables. All scripts for gene catalog construction and MAG reconstruction are available at the following GitHub repository (https://github.com/Minamehr/Manuscripts/tree/main/Dysbiosis-and-Functional-Shifts-IBS).

### Funding

Not applicable.

### Consent for publication

Not applicable

### Authors’ contributions

KBL and MHA designed the study. MHA and KBL conducted participant recruitment, clinical assessment, and sample collection. MHA developed the laboratory protocols. MHA, ME, and NKS performed the bioinformatics and statistical analyses. MHA drafted the manuscript with contributions from all co-authors. All authors reviewed and approved the final manuscript.

### Competing interests

The authors declare that they have no competing interests.

## Acknowledgment

The authors gratefully acknowledge the staff of the Gastroenterohepatology Research Center for their support during participant recruitment and sample collection. The authors acknowledge support by the state of Baden-Württemberg through bwHPC.

## Notes

### Competing Interest Statement

The authors have declared no competing interest.

