## Supplemental Data 2 for "Integrated gene-catalog and genome-resolved metagenomics reveal taxonomic and metabolic differences in the gut microbiome in irritable bowel syndrome"

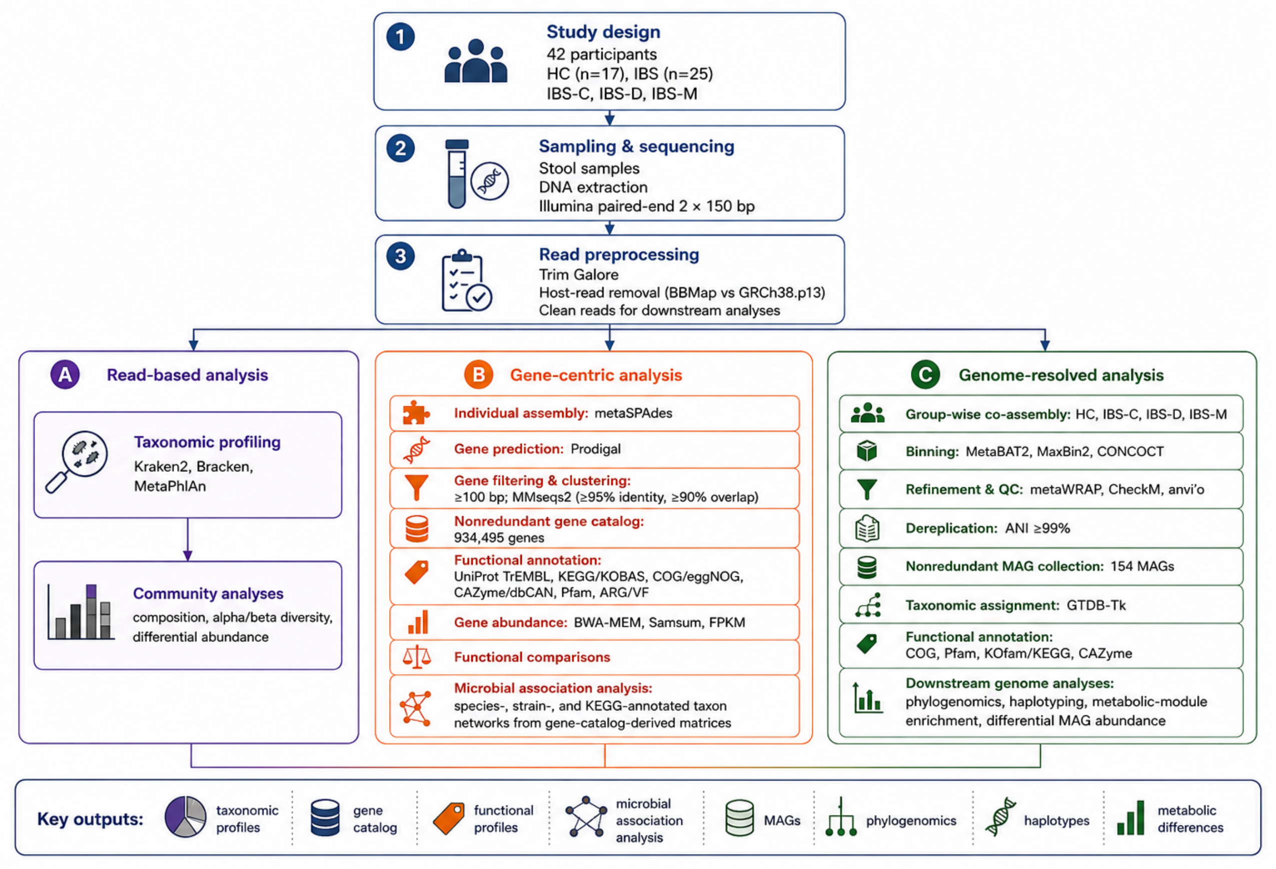


**Figure S1.** Overview of the metagenomic workflow used in the IBS study. Read preprocessing and read-based taxonomic profiling, gene-catalogue construction, and MAG reconstruction were performed using versioned tools and workflows within the Galaxy platform on the European Galaxy server (usegalaxy.eu). Downstream functional, genome-resolved, microbial association, and statistical analyses were performed using the indicated tools.

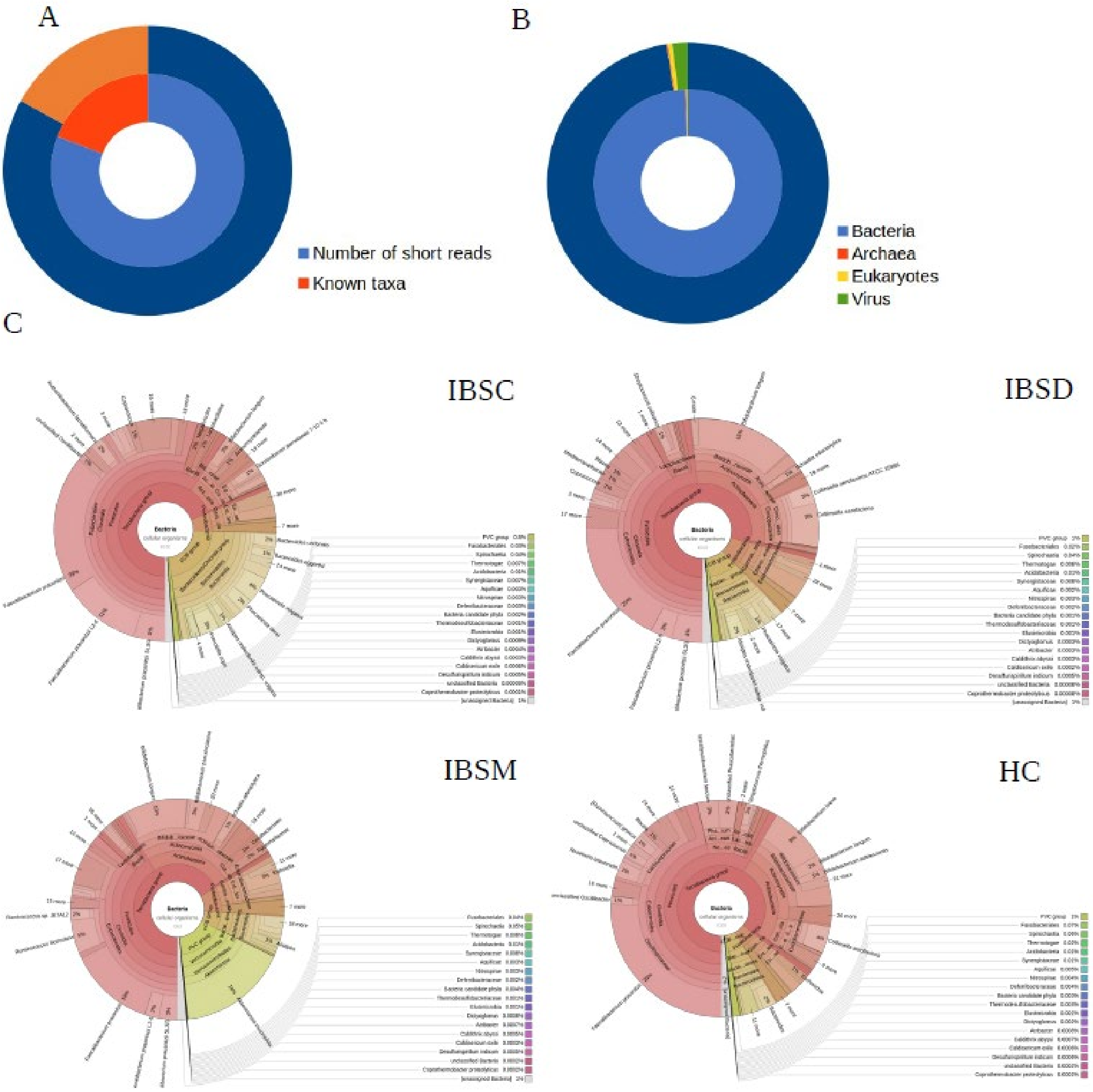


**Figure S2.** The taxonomic annotation of short reads using Kraken2. A: The numbers of high-quality short reads and known taxa (could be annotated by Kraken2) for patients with IBS and HC. B: The number of known taxa classified to different domains. C: Taxonomic composition of microbial communities from different subtypes of IBS (IBSC, IBSD, and IBSM) and healthy control (HC) estimated from high-quality short reads with KRAKEN2 and visualized with KronaTools.

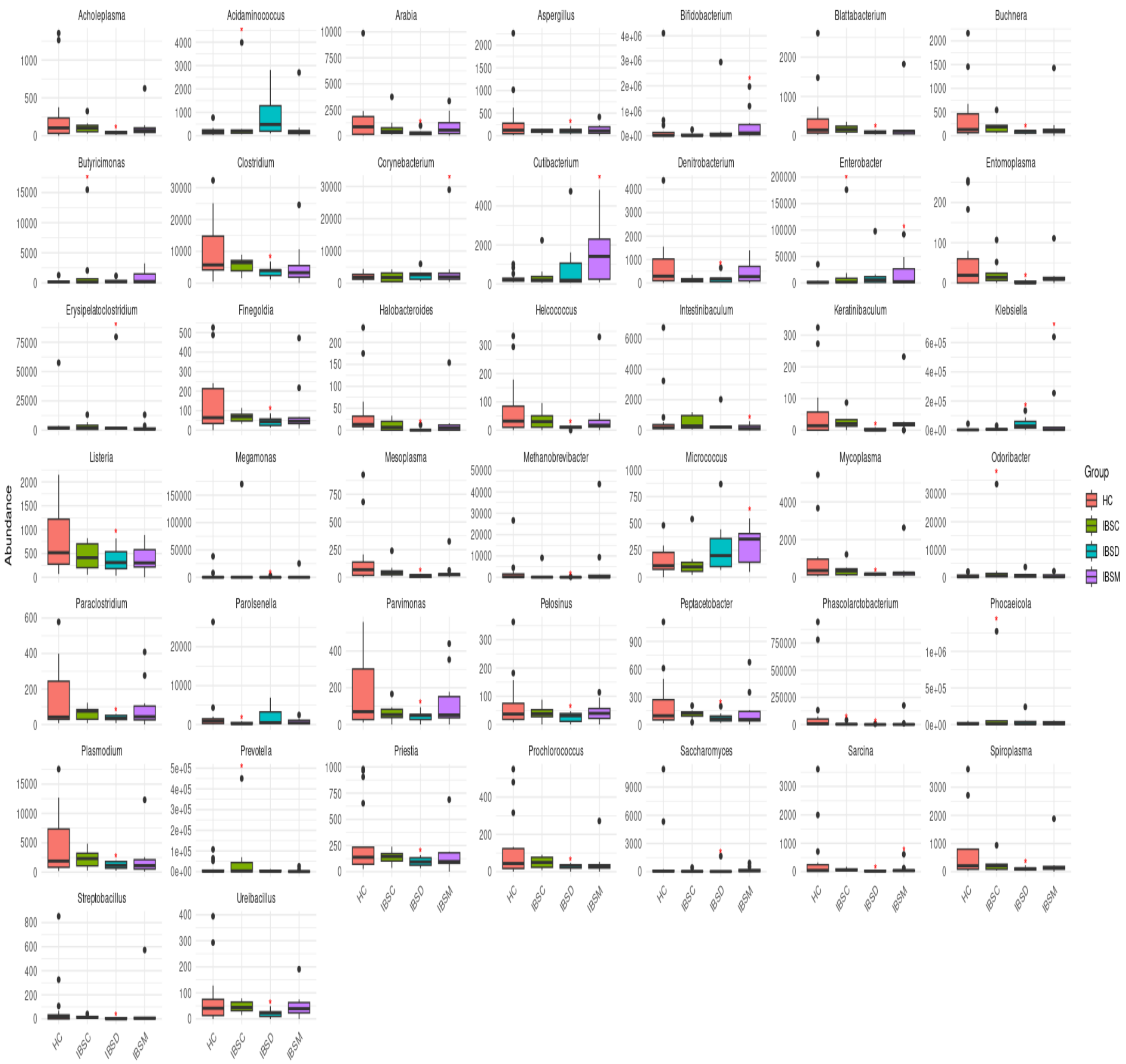


**Figure S3.** Differentially abundant genera between HC and IBS subtypes. Significant differences identified by DESeq2 are marked with a red asterisk (*). The colour codes for HC and IBS subtypes are shown in the legend. Differences in *Klebsiella* and *Clostridium were also supported* by MaAslin2.

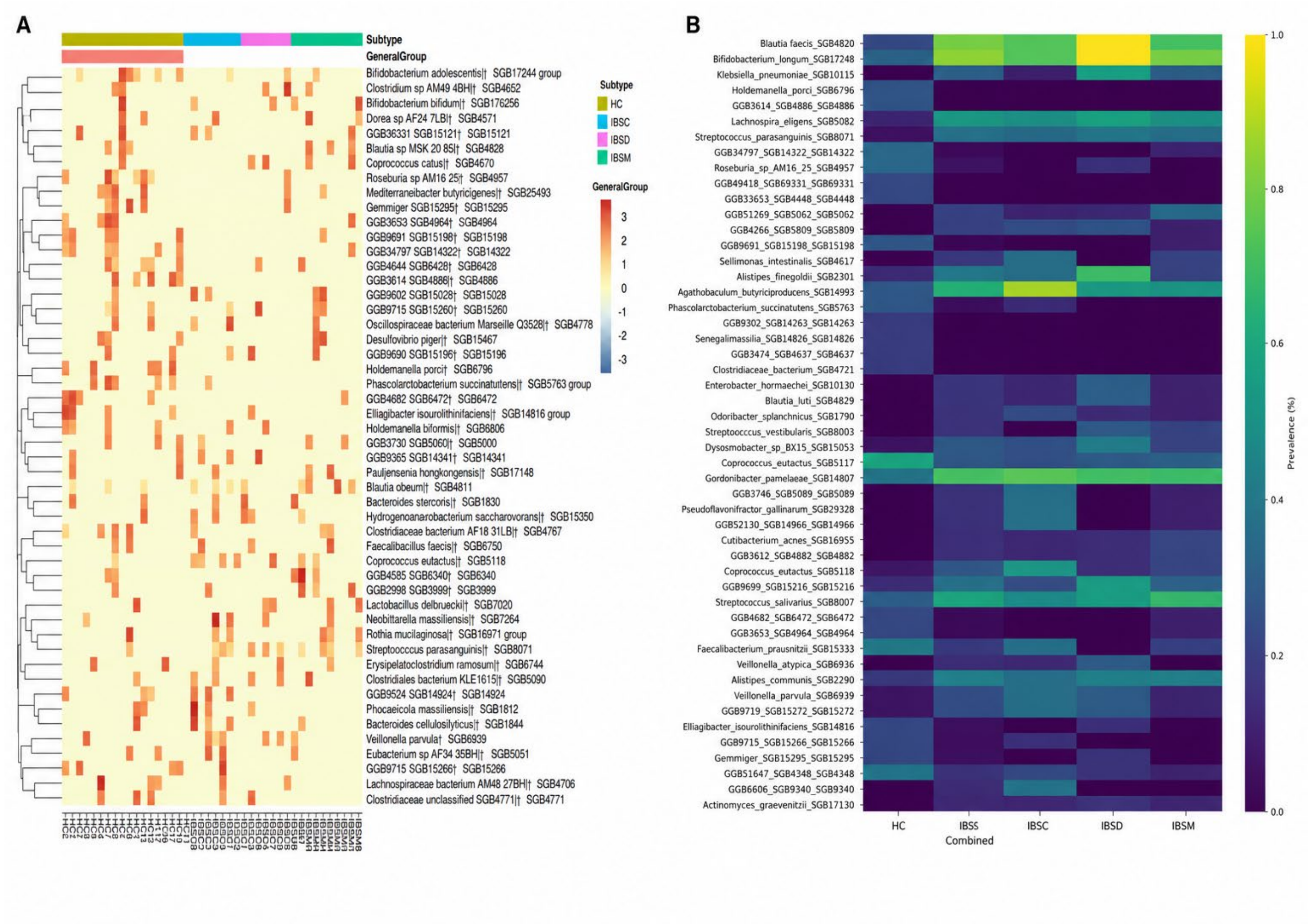


**Figure S4.** Differential abundances and prevalence of IBS-associated strains. A: The most significantly abundant strains differed between patients with IBS and HC. B: The most prevalent strains showed significant differences between IBS patients and HC.

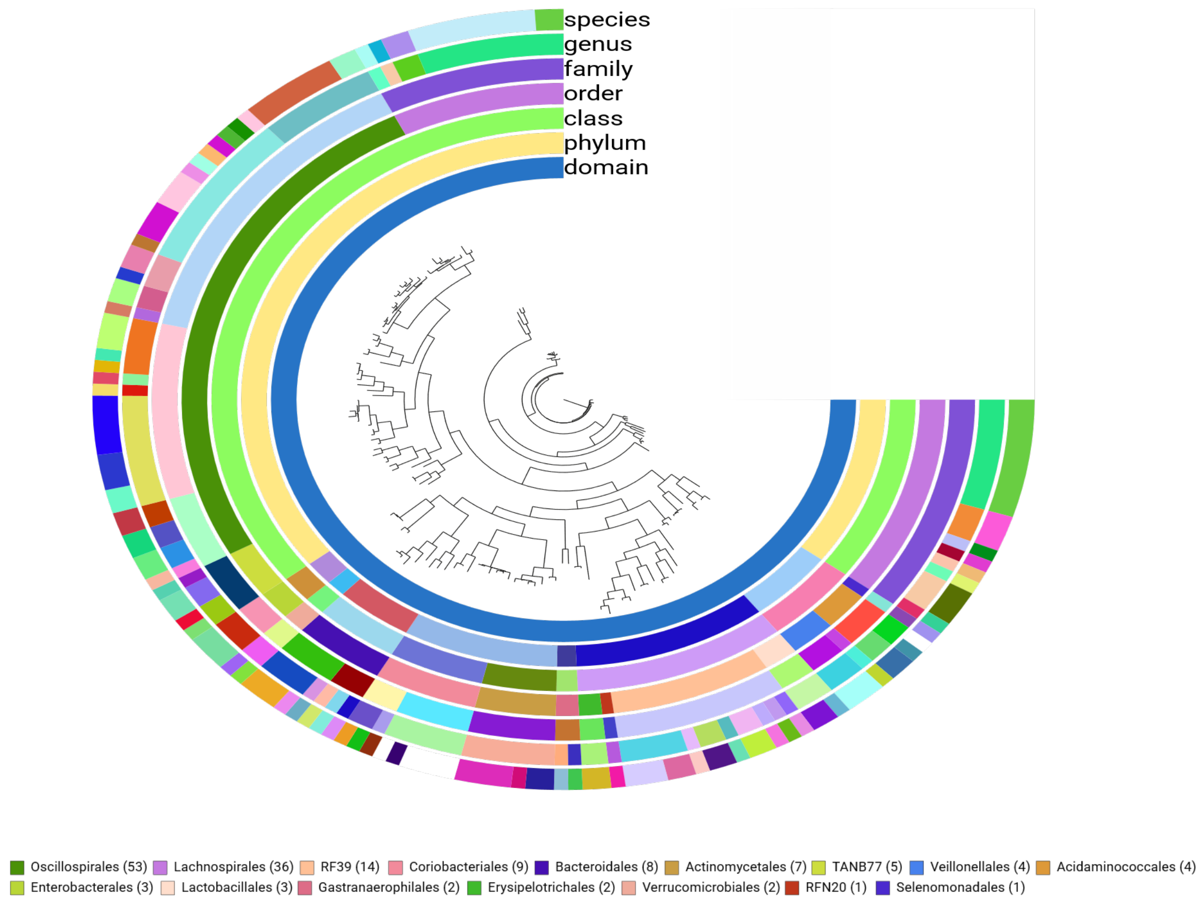


**Figure S5.** Genomic phylogenetic tree constructed using Anvi'o, based on 154 non-redundant MAGs. The tree was built from concatenated amino acid sequences of 37 single-copy ribosomal proteins using Anvi'o's phylogenomics workflow. Numbers preceding each taxonomic order label indicate the number of MAGs assigned to that order.

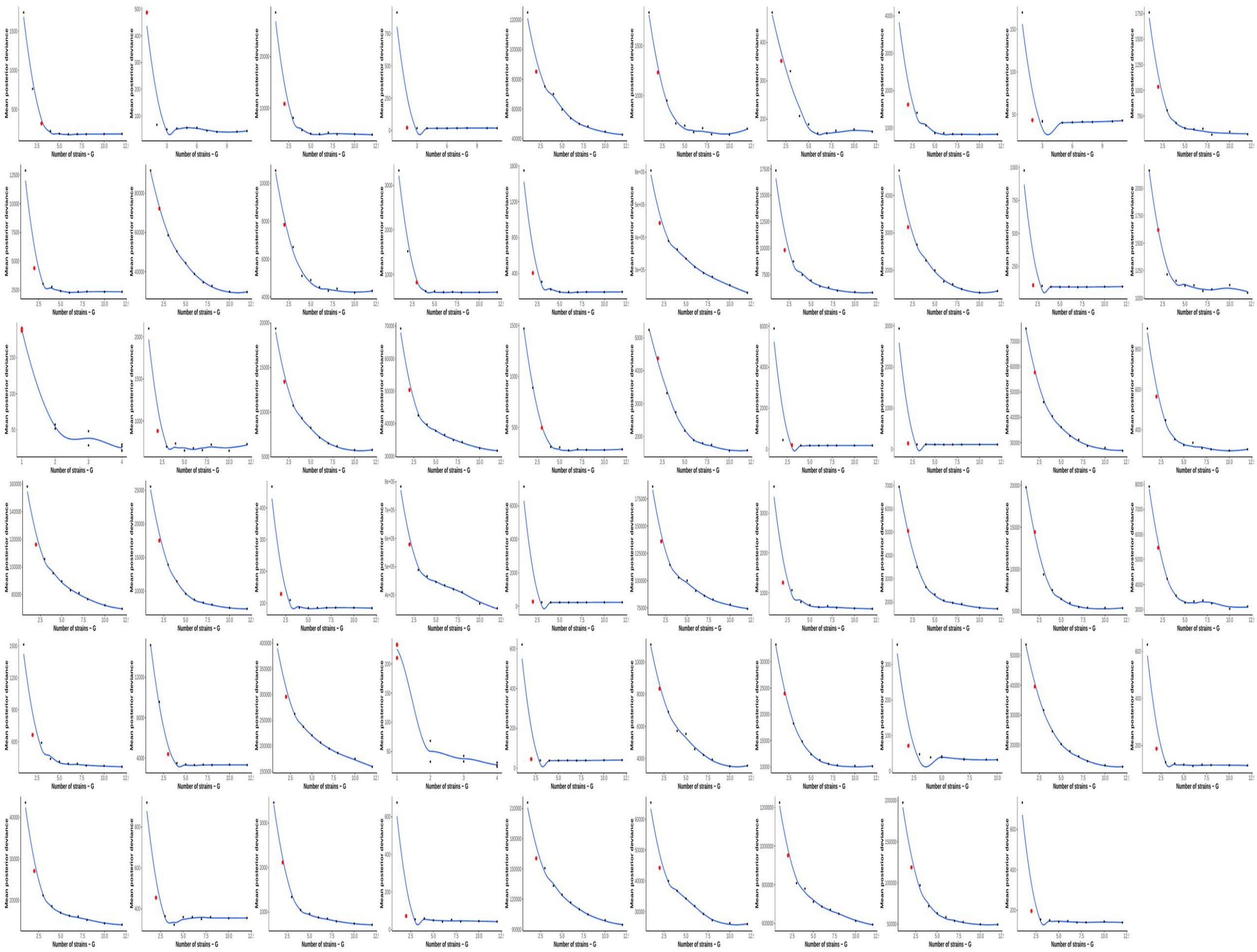

**Figure S6.** DESMAN haplotype decomposition for each MAG. Posterior deviance profiles across different numbers of inferred haplotypes (G) are shown for each of the non-redundant MAGs. The red dot indicates the estimated optimal number of strains, as determined by a heuristic based on posterior deviance reduction, haplotype abundance threshold, and prediction accuracy.
